# Testing the Limits of SMILES-based *De Novo* Molecular Generation with Curriculum and Deep Reinforcement Learning

**DOI:** 10.1101/2022.07.15.500218

**Authors:** Maranga Mokaya, Fergus Imrie, Willem P. van Hoorn, Aleksandra Kalisz, Anthony R. Bradley, Charlotte M. Deane

## Abstract

Deep reinforcement learning methods have been shown to be potentially powerful tools for *de novo* design. Recurrent neural network (RNN)-based techniques are the most widely used methods in this space. In this work, we examine the behaviour of RNN-based methods when there are few (or no) examples of molecules with the desired properties in the training data. We find that targeted molecular generation is often possible, but the diversity of generated molecules is often reduced, and it is not possible to control the composition of generated molecular sets. To help overcome these issues, we propose a new curriculum learning-inspired, recurrent Iterative Optimisation Procedure that enables the optimisation of generated molecules for seen and unseen molecular profiles and allows the user to control whether a molecular profile is explored or exploited. Using our method, we generate specific and diverse sets of molecules with up to 18 times more scaffolds than standard methods for the same sample size. However, our results also point to significant limitations of one-dimensional molecular representations as used in this space. We find that the success or failure of a given molecular optimisation problem depends on the choice of SMILES.

## 2 Introduction

Developing a novel drug is a complex and difficult problem.^1^ The pipeline from hit-to-lead, then through clinical trials, to market is plagued with failure.^2–4^ While there are several known challenges to improve the efficiency of the drug development process, one key step would be to produce better early hits by reducing resources spent on inappropriate molecules. Beyond hit generation, the ability to generate novel molecules with specific properties would be beneficial in much of the pipeline, improving cost, speed, and effectiveness.^5^

Ideally, during drug discovery, given a target and a required molecular profile, we would search for suitable molecules in all drug-like chemical space. Current experimental high-throughput methods do allow millions of molecules to be tested, however further increases in scale are still prohibitively expensive.^6^ Therefore, given that there are an estimated 10^60^ synthetically accessible drug-like molecules^7^, of which approximately 10^8^ have been synthesized^8^, experimental methods are not sufficient for comprehensive sampling of chemical space.

Computational methods offer the promise of being able to search larger areas of chemical space and virtual screening is commonly used to search curated chemical libraries for potential hits.^9–12^ However, the chemical space available for assessment is only a tiny proportion of the possible space.^13–15^

Instead of searching existing molecular datasets, computational *de novo* design models aim to create new sets of novel molecules.^12,16^ These generative models have become a popular choice in computational molecular design in recent years.^17-19^ Early *de novo* models were rules-based algorithms designed to generate molecules by enumerating fragment combinations.^17,18^ While these models were able to generate large sets of novel molecules, they required manual encoding of disconnection rules, filtering to generate chemically valid molecules, and often only enumerated fragments from available molecular datasets. Such early models were often paired with global optimisers to generate molecules with desired properties.^6,19,20^ Over the last few years, deep learning molecular generation tools have become more prevalent.^21^ Similar to early molecular generation algorithms, deep learning *de novo* tools often pair molecular generation with optimisation in order to produce focused sets of molecules with improved performance.

Several deep learning approaches have been applied to *de novo* molecular design. Autoencoders^22^ are perhaps the most frequently applied method for *de novo* molecular generation and optimization.^12,25^ Here the discrete representation of a molecule is converted to a continuous representation (encoded) from which its properties can be predicted and optimised. The resulting continuous representation is then converted back to a discrete molecular representation (decoded).^23,24^ Several popular *de novo* design methods use SMILES^23^ to discretely represent molecules, often with recurrent neural networks (RNNs). An early example of this approach used RNNs to generate a molecular library through a SMILES language model^16^, before fine-tuning the model on a smaller subset of molecules with desired properties. Another popular method to fine-tune SMILES generation models is reinforcement learning (RL). In theory, RL methods allow a user to generate a set of molecules with specific properties without explicit examples of molecules that match the reward profile.

In this paper, we examine on-policy reinforcement learning models.^25^ These models learn a policy to maximise the reward for each action taken. For molecular generation, the policy describes a strategy that maximises the reward for each generated molecule. This is done by augmenting a prior model to produce sets of molecules optimised for specific molecular profiles. In 2017, Olivecrona *et al*. proposed a policy-based reinforcement learning model, REINVENT^26^, to tune a SMILE-generating RNN. The authors reported that their model could generate novel molecules similar to a target structure even after analogues of the target were removed from the training data. Furthermore, it was able to generate predicted actives (predicted activities greater than 95%) to the DRD2 receptor even when known actives were removed from training. Popova *et al*. proposed another using the same general architecture (ReLeaSE) that was able to bias the generated molecules against structural complexity, melting point, and activity against the JAK2 protein. In 2022, the authors of REINVENT proposed an update to their method that enables the use of curriculum learning to fulfil complex generation tasks while reducing the overall cost of learning.^27^ These recent papers point to the potential of SMILES-based deep RL models in the generation and optimisation of novel molecules. However, it is still not clear how the prevalence of desired molecular profiles in training data will affect optimisation performance.

In drug discovery, successful hit generation requires new molecules with novel combinations of properties. These molecules often do not exist in currently available datasets; therefore, for RL methods to be truly useful, they must be able to extrapolate. At present, it is unclear how well these models are able to do so. It is also important that RL methods can generate sets of molecules that explore the chemical space for a complex molecular profile and produce a diverse library and exploit the chemical space to generate focused molecular libraries. In this paper, we explore scenarios where optimisation is attempted with little or no representation in the training data and investigate the extent to which the methods can extrapolate. We manipulate the prevalence of specific properties, measured as a percentage of the entire training dataset, and test the limits of optimisation of individual properties. We find that the RL models tested can extrapolate beyond the training data but often produce a set of molecules with little diversity. We show that these models are frequently unable to generate molecules that satisfy complex molecular profiles. We go on to demonstrate a curriculum-learning-inspired optimisation procedure that enables the generation of specific and diverse sets of molecules that satisfy complex and unseen molecular profiles. We also highlight the limitations of SMILES-based molecular generation tools.

## 3 Methods

We assessed the performance of a popular on-policy SMILES generation model, REINVENT^26^, to determine the limits of deep RL tools in molecular design. Like earlier RL molecular generation tools^28^, REINVENT involves a two-step process. The first is to train a prior RNN to generate SMILES through supervised learning. This model is trained to correctly predict the next character of a SMILES string given a starting token or incomplete string. The second is to fine-tune the prior model producing an agent model able to generate a focused library through a reward-feedback loop. During this second step, the model learns a policy that maximises the likelihood of generating a molecule with a favourable reward (Figure 1). Full details of the models can be found in the work of Olivecrona *et al*.^26^

**Figure 1:**
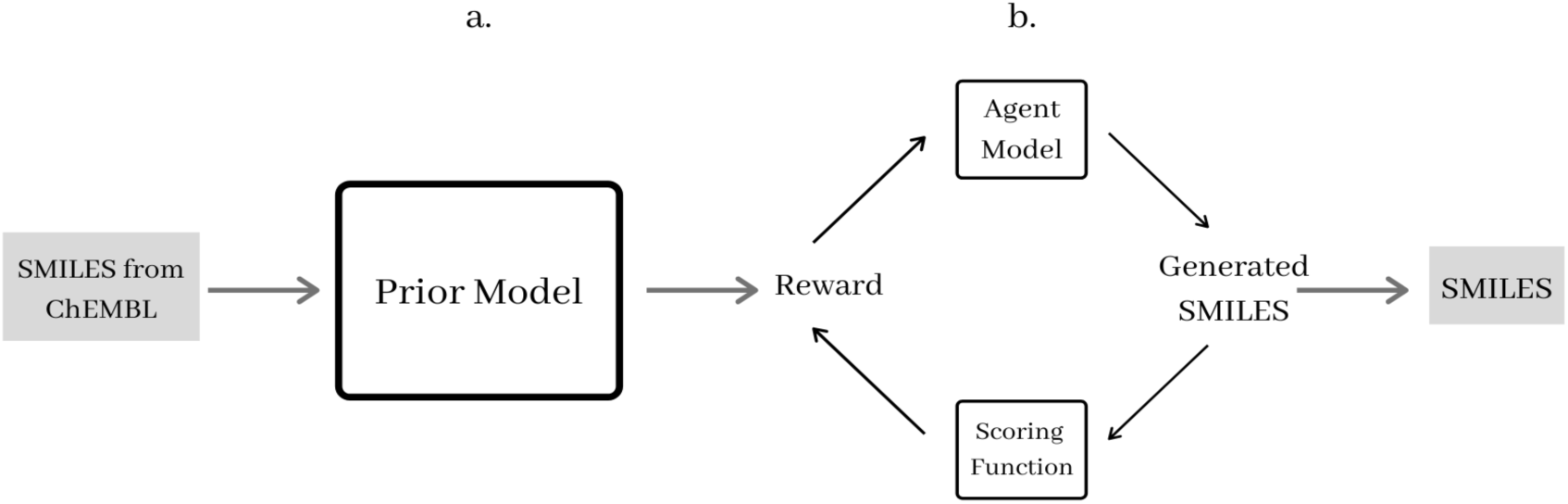
High-level diagram displaying the architecture of the deep reinforcement learning model used by Popova et al and Olivecrona et al. ^29, 30^ (a) Supervised learning language model. The prior model learn s to generate novel SMILES from a large dataset of SMILES from ChEMBL. ^31^ (b) Reinforcement Learning. The agent model (based on the prior model) is trained to generate SMILES that return a favourable reward.

For all experiments described in the following, the model was trained on subsets of 1.5 million drug-like molecules from ChEMBL^29^. After the model was fully trained, we sampled 500 molecules unless otherwise stated. We chose to generate 500 molecules as this provided a large enough sample from which we could draw clear conclusions about the distribution of generated molecules.

### 3.1 Property Characterisation

Hydrogen bond acceptors (HBA), donors (HBD), molecular weight (MW), topological polar surface (TPSA) and LogP were calculated using the chemical descriptor module from RDKIT.^30^ Synthetic accessibility (SA)^31^, QED^35^, and Tanimoto similarities were also calculated using RDKit.

For the analysis of generated molecular sets, only unique molecules (all valid SMILES generated once repeats are removed) were considered. To compare the diversity of training and generated datasets, Murcko scaffolds^32^ were generated using RDKit. The generated library internal diversity scores were calculated using MOSES.^33^

### 3.2 Reinforcement Learning

To successfully optimise for a property, a suitable reward function must be provided. For simplicity, throughout our study, we used the same step reward function (examples available in the supplementary information). Any invalid SMILES did not return a reward and all valid SMILES that met the reward criteria returned a reward of one.

#### 3.2.1 Optimisation Success

We initially determined the success of optimisation using the proportion of molecules in the generated library that fell within the reward range. However, after several properties were tested at increasing representations, it became clear that the difference between the proportion of optimised molecules in successful and unsuccessful optimisation attempts was large enough so that a success threshold was not appropriate. Instead, optimisation attempts that showed an increase in the proportion of optimised molecules were deemed successful. This was possible because all unsuccessful attempts resulted in zero optimised molecules.

### 3.3 Recurrent Iterative Optimisation Procedure

We propose a novel, curriculum learning inspired recurrent Iterative Optimisation Procedure (rIOP). Curriculum learning is a method used to teach models how to complete difficult tasks through the gradual introduction of more complex examples during training.^34^ For single-step optimisation attempts, it is common for RL methods to exploit molecular motifs found to return positive rewards, leading to generated sets with low diversity (specialisation). We expect that the greater the difference between the reward profile and training data, the more prevalent this behaviour is. By splitting the optimisation task into a series of smaller tasks, we reduced the difference between the molecules generated by the prior and the desired reward profile at each step. Thus, reducing the likelihood of early specialisation. Repeating a prior/agent training loop with a series of small changes in the reward profile, we encouraged each agent model to shift its property distribution toward the final, desired, distribution. The resulting agent was then used as the prior in the next step. Splitting the final optimisation task into a series of increasingly complex subtasks allowed the model to satisfy increasingly difficult molecular profiles that directed it toward the final goal.

We demonstrate the use of two implementations of rIOP. The first, single model rIOP (SrIOP), only samples from the previous agent when training the current model. The second, double model (DrIOP), samples from the previous two models. Unless otherwise stated, for DrIOP we sampled the current agent once for every five times the previous agent was sampled.

#### 3.3.1 Diversity filters

To control diversity, where appropriate, we incorporated the diversity filters described by Blaschke *et al*.^35^ With diversity filters (DF) enabled, the model will only give a positive reward for the first *n* molecules that satisfy the reward function for a given scaffold. Once *n* molecules that match the reward profile have been generated, molecules with this scaffold are no longer rewarded. This prevents the model from entering a local optimisation minimum by producing many molecules with the same scaffold and small structural differences to satisfy the reward function.

## 4 Results

Deep RL molecular generation models are powerful tools for optimising molecular properties. However, their usefulness is dependent on their input training data. We show how these tools are able to optimise for specific property values, however, only within a property specific value range. We also show how these methods can generate molecular profiles that are not present in training data and how the representation of training data affects the composition of generated sets of molecules.

By evaluating the performance of deep RL molecular generation methods with increasing proportions of training data that match a desired property profile, we show the effects of representation on generated molecules. To overcome the generated library restrictions caused by training data representation, we propose a curriculum learning inspired approach (rIOP) that allows for the optimisation of under-represented properties. rIOP also allows users to optimise generated molecules toward complex molecular profiles that are not possible with REINVENT while controlling the diversity of generated libraries.

### 4.1 Control of Generated Libraries

A standard goal for *de novo* design deep RL tools is to produce novel molecules for which the property distribution of a single property or many properties has been controlled. Previous studies using methods such as REINVENT and ReLeaSE have shown that it is possible to bias generated molecules toward specific properties such as hydrophobicity, melting point, or predicted activity against the DRD2 receptor.^26,28^

To determine the degree to which the property distribution of the molecules generated by the agent (agent distribution) can be controlled, we set REINVENT the task of shifting the distribution of LogP or HBA counts across their respective ranges. These properties may not be the most important in a drug discovery context; however, these experiments allow us to assess optimisation performance. If it is possible to control the distribution of generated molecular sets, we expect to observe changes in the composition of these sets as the reward range changes. During RL, all valid SMILES that satisfy the reward function are given a reward of one. All other molecules, valid or not, receive zero reward.

Figure 2a shows the property distribution of generated molecules for the reward ranges of LogP between -15 and 20. It shows that it is possible to control the position of agent distributions with the reward range. However, in extreme cases (LogP reward between -15 and -10), optimisation was unsuccessful, and we observe no change in the agent distribution; the training data distribution is reproduced.

**Figure 2:**
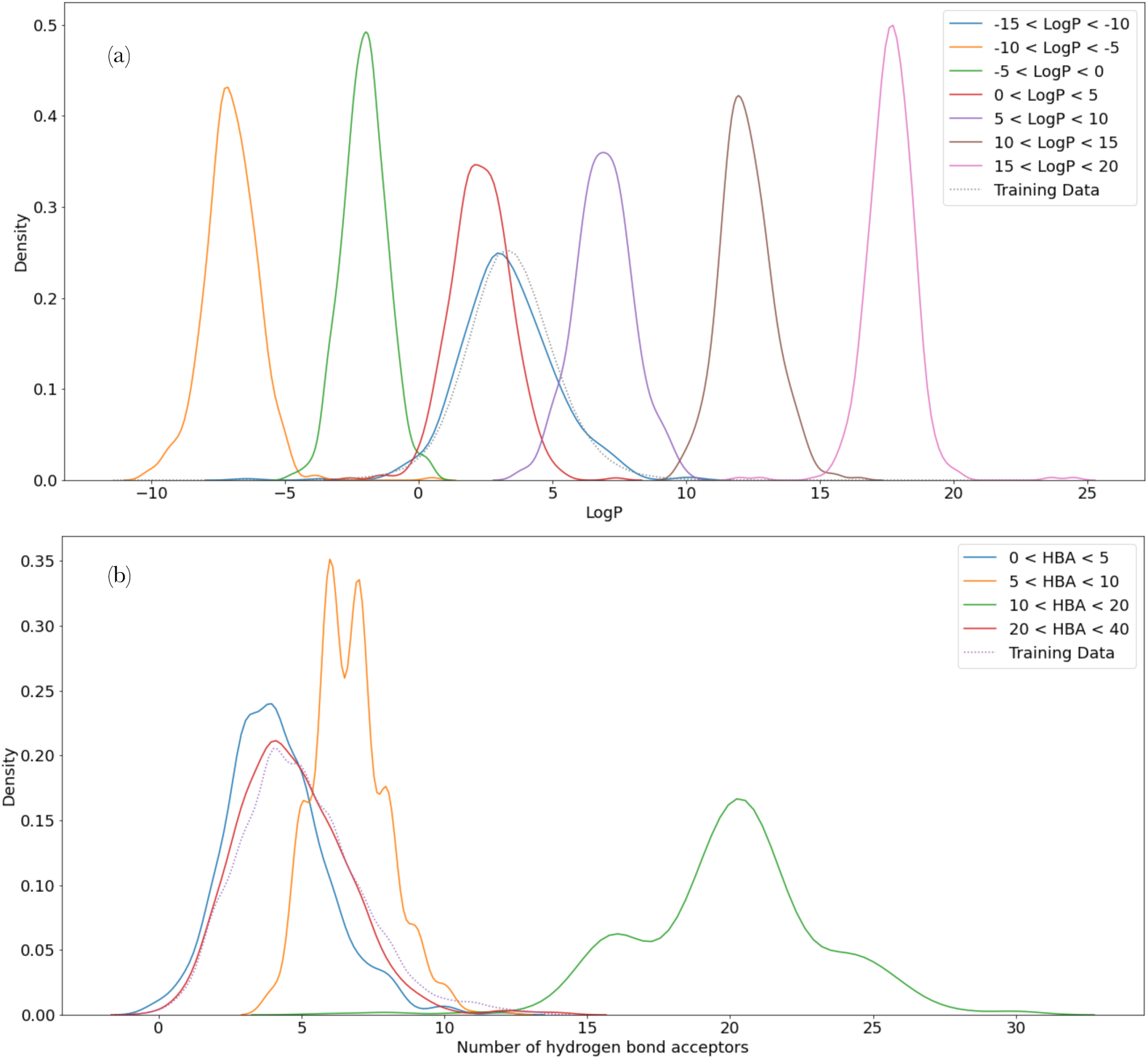
Distribution of generated libraries for (a) LogP optimisation and (b) number of HBA’s optimisation. Each line corresponds to the property distribution of the molecules sampled by an agent trained with a reward range detailed in the legend. Both figures show that optimisation is possible within a specific range (e. g., 0 - 20 HBA’ s), outside this range optimisation fails. The model is unable to generate appropriate molecules, so the training data distribution is recreated.

The same behaviour was observed for HBA counts (Figure 2b). Optimisation anywhere in the range of 0-20 was possible; however, optimising the model to generate molecules with 20 HBA’s or more was unsuccessful (red distribution). As with LogP, a distribution similar to the training data was reproduced. The training data did include several molecules with more than 30 HBA’s. A full breakdown of the generated molecular sets, reward ranges, and examples of molecules can be found in the supplementary information.

We postulate that this failure occurs because in the extreme case fewer molecules generated by the prior model return any reward during RL. If trained for infinite time, the model will eventually randomly generate SMILES that will return a positive reward; nevertheless, poor representation can prevent effective optimisation. This ineffective optimisation then leads to the model repeatedly producing the same SMILES seen in training; thus, the training data is reproduced.

### 4.2 Effect of percentage representation

We have shown that using deep RL molecular generation tools, optimisation of under-represented properties is sometimes not possible. To investigate how widespread this issue is, we tested the ability of the models to generate molecules with properties within and outside the training data.

We prepared several training datasets in which the proportion of molecules that matched the desired reward profile varied. We chose reward profiles (supplementary information) at the upper end of the full ChEMBL training data distribution such that at least 10% of the training data matched the reward function. Once the reward range was calculated, all molecules that matched the reward profile were removed from the full training data set. Then smaller random samples equal in size to 0%, 2%, 5%, 7% and 10% of the entire training dataset were put back and used to train the model from scratch.

Table 1 shows that using REINVENT optimisation was successful for all properties across all percentage representations. In all cases, the generated set distributions were shifted toward our desired property profile relative to the training data. These results show that, for the properties tested, the model was able to learn the chemical-structural relationship from the surrounding molecules in the distribution; it is possible to learn without representation in the training data.

**Table 1:**
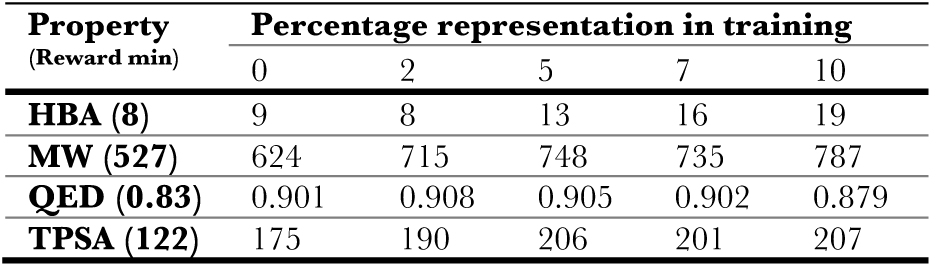
Mean property values for molecules generated from four property optimisation tasks using increasing training data percentage representations.

Table 1 shows that the composition of each generated library is dependent on the representation of the desired property in the training data. For example, we can generate sets where most molecules have a HBA count greater than 8 (the reward threshold), but a higher percentage representation leads to molecules with higher HBA counts with the same reward function. The mean of the 0% representation is 9 HBA’s compared to 20 for the 10% experiment. Directional changes across each experiment can be seen for all properties tested (supplementary information). We postulate that the trends in the generated molecules mirror the training data. For example, for a specific reward property value (e.g. HBA = 5) if, as the percentage representation of that property profile increases, the diversity of the training data increases, you will see an increase in the diversity of the generated library. Conversely, if all the examples in the training data were very similar, you would observe a reduction in training data diversity and generated library diversity. Therefore, for most properties, we expect that a lower percentage representation would lead to a less diverse generated library.

These results show that it is possible to generate molecular profiles that are not seen during training and that the composition of the generated molecular sets depends on the degree to which the desired molecular profile is prevalent in the training data. Therefore, depending on the use case for these models, different percentage representations for training may be suitable. However, when the aim is to generate an unseen molecular profile (0% representation), standard methods leave the user without control over the composition of the generated library.

### 4.3 Curriculum Learning for Generated Library Control

We have shown that it is possible to generate molecules with unseen molecular profiles in training data (Table 1). However, the model’s ability to do this is limited at the extremes of the property distribution (Figure 2). The composition of each generated set depends on the prevalence of molecules in the training data with the desired molecular profile. Building on previous work^27^, we postulate that higher training data representations would often lead to greater diversity for generated molecules, as the model would often have a more diverse set of examples to learn chemical-structural relationships.

To improve the efficacy of deep RL generation methods, we propose a new curriculum learning-inspired approach, called recurrent iterative optimization procedure (rIOP). Our method allows deep RL generation methods to maximise the diversity of generated molecules for seen and unseen molecular profiles during optimisation. It also enables the model to generate molecules that perform more complex optimisation tasks where standard methods fail.

#### 4.3.1 Recurrent Iterative Optimization Procedure to Improve Diversity

To demonstrate how rIOP can increase the diversity of simple optimisation tasks, we generated molecules with a reward for TPSA between 250 and 300 using SrIOP and REINVENT’s standard implementation. Standard methods do generate molecules in this range; however, we expect that with SrIOP we will see an improvement in the diversity of generated molecules. TPSA shift is a simple optimisation task; therefore, we only sampled the agent of the previous step when training the current prior (see methods).

Figure 3 shows each step of the SrIOP procedure and the change in property distribution at each stage to match the reward function. To measure the diversity of each generated library, we calculated the number of unique Murcko scaffolds generated. With each SrIOP step, we see a reduction in the number of scaffolds generated. This is to be expected as we are moving toward the limit of the full TPSA property distribution, where there are fewer ways to achieve these property values. Table 2 shows that in our last step (SrIOP 4) we produce 18 times more scaffolds using SrIOP (55 scaffolds) compared to REINVENT (3 scaffolds). Furthermore, SrIOP generates more molecules from the 500 sampled that match the reward profile (494 compared to 476 for SrIOP and REINVENT, respectively). Examples of generated molecules can be found in the supplementary information.

**Figure 3:**
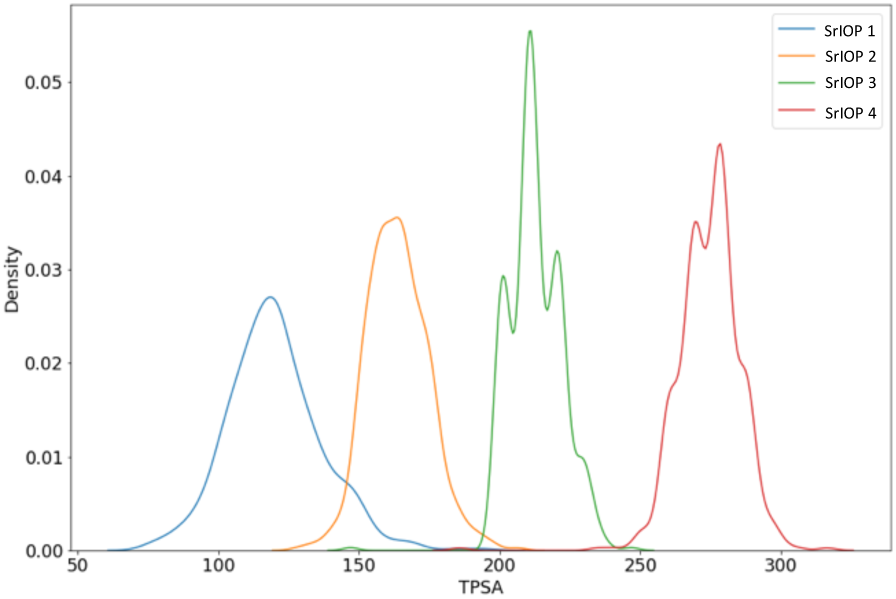
TPSA distribution of molecules sampled from each intermediate (1 - 3) and final (4) agent trained during rIOP. We show that we can shift the distribution iteratively towards target property values. TPSA reward range for each step were (1) 100 – 150, (2) 150 - 200, (3) 200 - 250, (4) 250 - 300.

**Table 2:** Comparison of generated set composition of each agent trained during SrIOP and REINVENT for TPSA optimisation.

| Run | # In reward range | # Scaffolds |
| --- | --- | --- |
| **SrIOP 1** | 437 | 352 |
| **SrIOP 2** | 468 | 198 |
| **SrIOP 3** | 499 | 20 |
| **SrIOP 4** | 494 | 55 |
| **REINVENT** | 476 | 3 |

#### 4.3.2 Iterative optimisation procedure for diversity control

For *de novo* design tools to be effective, it should be possible to control the specificity of the generated molecules. We have shown how the representation of a desired profile during training can affect the composition, and hence the specificity, of generated sets of molecules. Our results also show how the use of our new method, SrIOP, can improve the diversity of molecules generated during simple optimisation tasks.

To demonstrate how SrIOP can be used to control the diversity of generated molecules, we created a library of compounds where our aim was to maximise QED. We use no molecules with QED greater than 0.8 during training, then iteratively increased the QED reward profile at each step. Diversity filters (DF) (see methods) were also used to further improve the diversity of generated molecules. DF prevents the model from producing the same scaffold repeatedly to maximise the reward. As in the previous experiment (Section 4.3.1), we only sampled the agent from the previous step when training the current agent (SrIOP).

Figure 4 shows how with and without diversity filters it is possible to generate molecules with high QED values. Figure 4b also shows that a wider distribution of molecules is produced with a diversity filter enabled. Of the 500 molecules sampled, SrIOP generates 490 molecules that match the reward function compared to only 317 using REINVENT. Of the 490 molecules, SrIOP generated 22 scaffolds. Standard methods produced more with 130 scaffolds across those 317 molecules. However, with DF enabled, we observed a significant increase in SrIOP performance, with scaffold diversity increasing from 22 to 297 in the 301 generated molecules. We also see a change using REINVENT; the model generates 234 scaffolds across 236 molecules. These results show how, using our SrIOP, we can generate a specific library of molecules (without DF), and a diverse library with DF enabled. On the contrary, REINVENT is only able to generate diverse molecules, eliminating the user’s ability to control the composition of generated sets. A full breakdown of the results, including each step of SrIOP, is available in the supplementary information.

**Figure 4:**
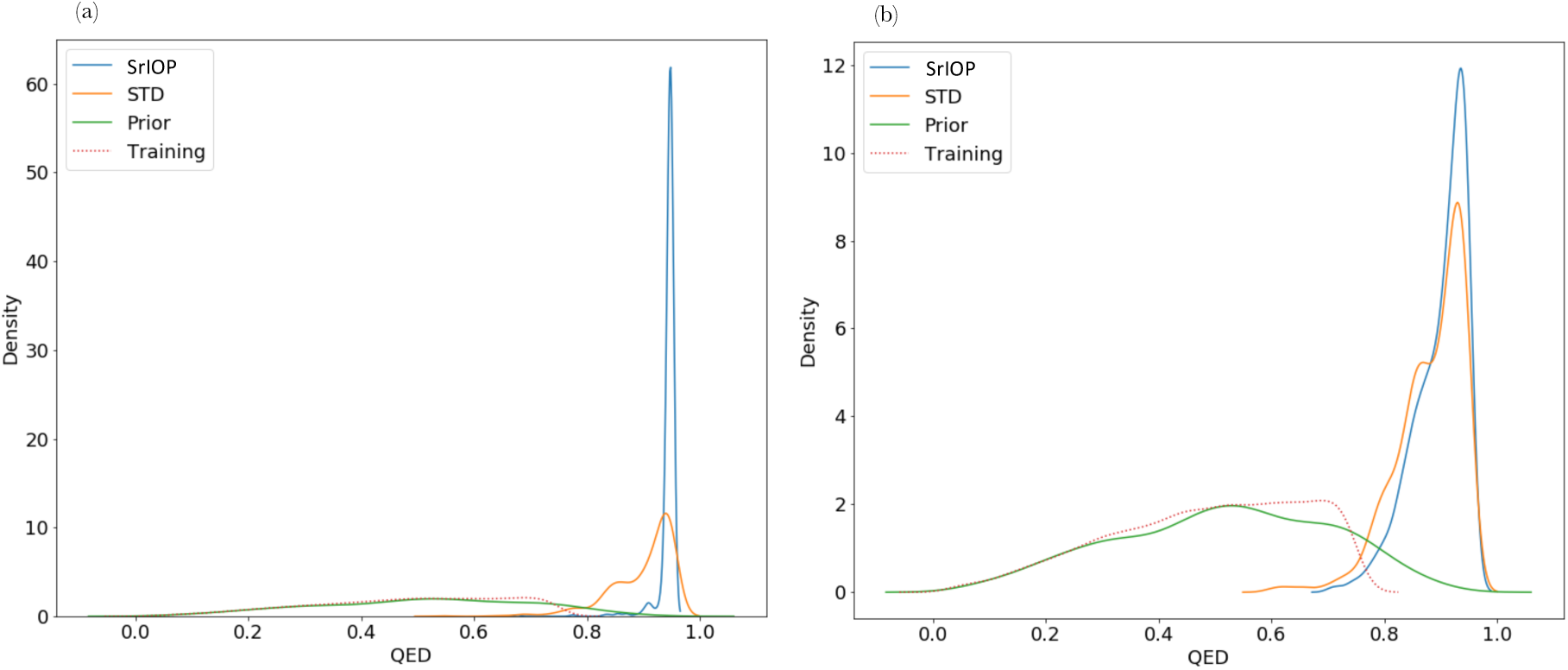
Comparison of SrIOP to REINVENT (STD) for generation of druglike molecules (a) without diversity filters, (b) with diversity filters. SrIOP (blue), standard (orange), prior (green) and training data (dotted) distributions for the generation of high QED molecules (QED > 0. 90). Only low QED molecules (< 0. 8) were used in training, then the QED reward range was increased each SrIOP step. Both methods can generate the desired molecules, however, SrIOP generates more molecules in the desired range in both cases.

SrIOP gives the user more control over the specificity of the generated library. If they wish to exploit a property, SrIOP used without DF filters will return a very narrow selection of molecules that match your reward profile. On the contrary, if a diverse library is required, enabling diversity filters with SrIOP will produce one. In this example, we chose a simple optimisation task as there are many ways to increase QED for a molecule. Therefore, we expected that the standard method would perform well. SrIOP still outperforms REINVENT in both specific (no diversity filter) and diverse (diversity filter used) set generation; however, we expect the difference in performance to increase for more complex optimisation tasks.

### 4.4 Comparison to other curriculum learning methods

Curriculum learning has long been used as a tool to overcome complex machine learning problems in various applications.^36^ However, its use in deep RL molecular generation tools is limited. There is one implementation of a similar method by the original authors of REINVENT^27^, that applies a curriculum learning approach to solve complex optimisation tasks, which we call ReCL.

#### 4.4.1 Iterative Optimisation procedure for complex optimisation tasks

One common use case of deep learning RL models is to optimise for molecules similar to a target structure. In such scenarios there may be few examples of the target structure in training data. To determine how useful SrIOP is in this practical situation, we have used it to generate molecules similar to target structures (Figure 5) with increasing difficulty.

**Figure 5:**
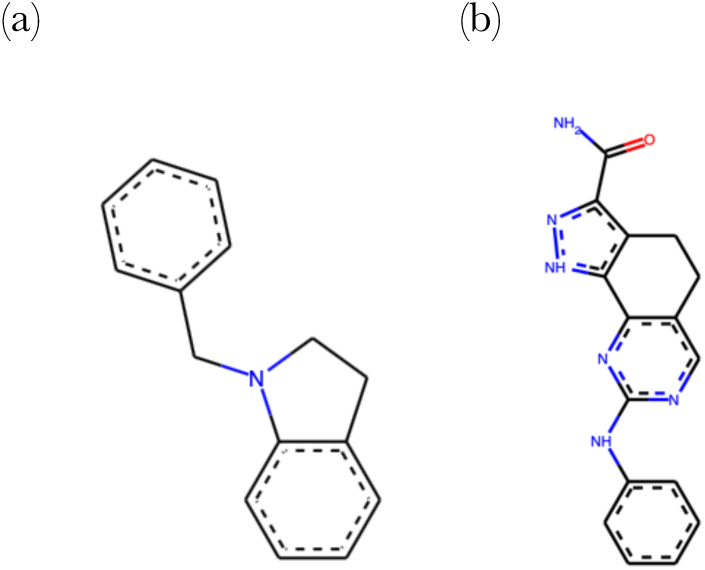
Diagrams of (a) simple and (b) complex target structures used for tanimoto similarity and substructure generation experiments.

In our experiments, we removed all molecules with tanimoto similarity greater than 0.4 to each target from the training data and then increased the tanimoto similarity reward threshold by 0.1 each step. Figure 6a shows how for a simple molecule (Figure 5a) it is possible to generate molecules identical to the target. Both SrIOP and ReCL perform well, with SrIOP generating more molecules with a tanimoto similarity between 0.9 and 1.0 (486 and 325 for SrIOP and ReCL, respectively).

**Figure 6:**
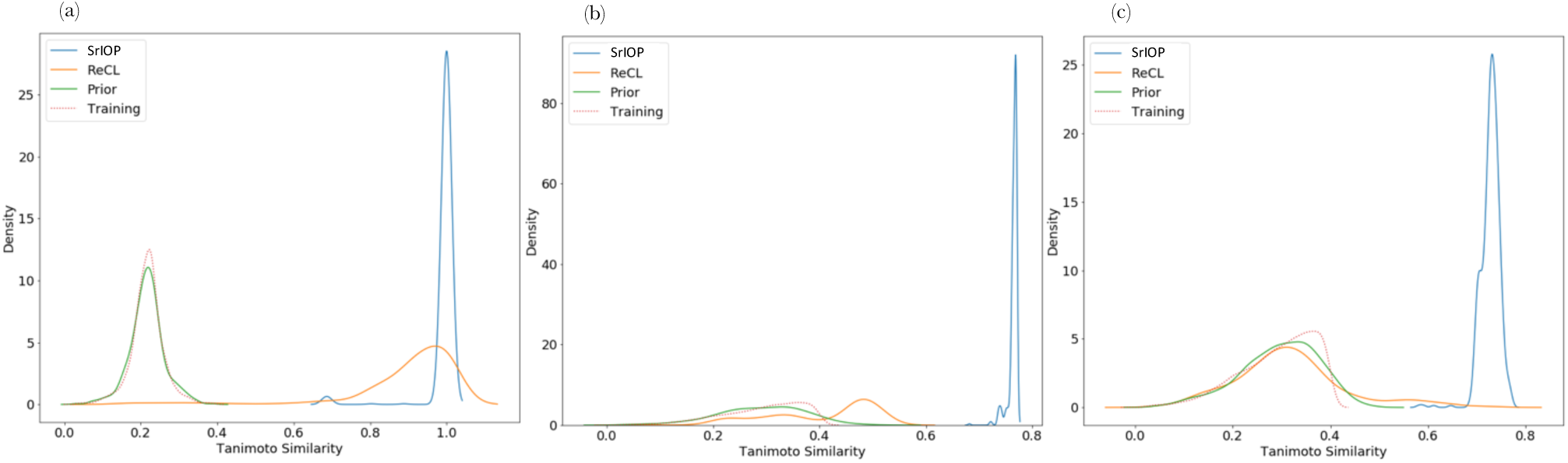
Generating molecules similar to (a) a simple target molecule (Figure 5 a), (b) a complex molecule (Figure 5 b) without diversity filters and (c) a complex molecule with diversity filters. SrIOP (blue), ReCL (orange), training (dotted) and Prior (green) distributions for each optimisation task. Both methods can produce entire datasets that match the simple target structure, however, only SrIOP is able to generate molecules similar to the complex target structure.

For a more complex molecule (Figure 5b), ReCL is less successful (Figure 6b and Figure 6c) as it cannot generate any molecules with a high similarity to the target structure (tanimoto similarity greater than 0.7). For SrIOP almost all (497 of 500 sampled) generated molecules have a high similarity to the target structure.

A second benefit of our method is the ability to control the diversity of the generated library.

Table 3 shows how, without a diversity filter, all 497 molecules sampled have the same scaffold. However, with diversity filters enabled, SrIOP generates 142 scaffolds across 447 molecules. In this example, we highlight the ability of SrIOP to fulfil more complex reward functions where similar CL methods fail. We show how it can also be used to control the diversity of the generated library through the inclusion of diversity filters.

**Table 3:** Breakdown of generated sets using SrIOP and ReCL in target similarity optimisation for (a) simple molecules and (b) complex molecules. For simple molecules, (a), a high similarity was all molecules with tanimoto similarity greater than 0. 9. For complex molecules, (b), the threshold was a tanimoto similarity greater than 0. 7. Diversity filters (DF) were used on the complex molecule.

| Method | # High similarity | # Scaffolds |
| --- | --- | --- |
| **SrIOP** | 486 | 1 |
| **ReCL** | 448 | 2 |

(b)
| Method | # High similarity | # Scaffolds |
| --- | --- | --- |
| **SrIOP** | 497 | 1 |
| **SrIOP (DF)** | 447 | 142 |
| **ReCL** | 0 | 0 |
| **ReCL (DF)** | 3 | 3 |

#### 4.4.2 Generating Complex Substructures

In their paper, Guo *et al*.^27^, report how they use ReCL to generate a library of molecules that contain the dihydro-pyrazoloquinazoline scaffold (Figure 5b). They do this by training their agent to generate molecules containing a series of increasingly complex substructures that direct the agent toward the final task. To compare our methods, we carried out the same experiment. Figure 7 shows each substructure and how, with our DrIOP method (blue) sampling from the two previous agents, we are also able to generate each substructure. Using DrIOP, we generated 103 molecules (from the 500 sampled) that included the final substructure across 21 distinct scaffolds. ReCL was able to generate more molecules with 451 molecules across 94 distinct scaffolds. This result suggests that DrIOP can generate molecules with increasingly complex substructures.

**Figure 7:**
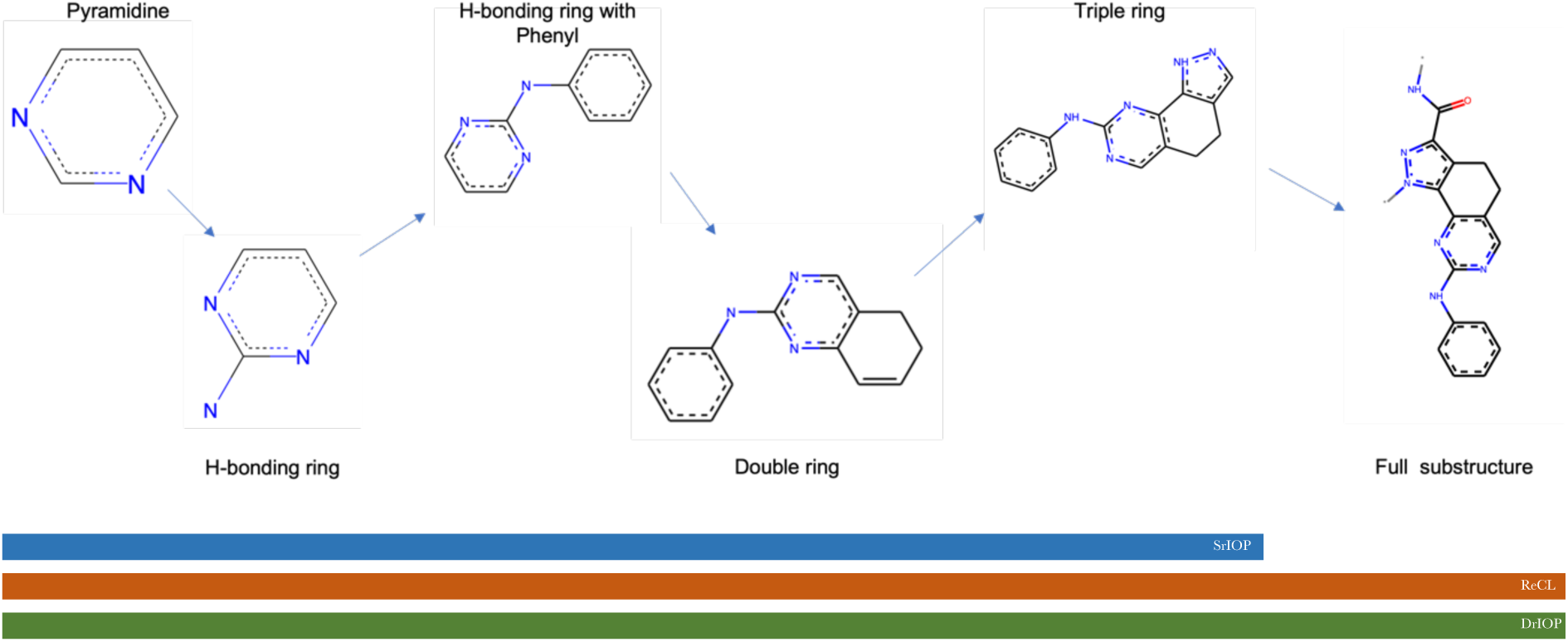
Substructure tree used to generate complex unseen molecules. Each method a ttempts to generate each substructure before moving on. SrIOP (blue) can only generate molecules with the first five substructures. ReCL (orange) DrIOP (green) can generate all six.

#### 4.4.3 Effects of Sampling from Multiple Agents

We have used rIOP where the previous one or two agents were sampled during reinforcement learning. For simple optimisation tasks, we used only the previous agent. However, for more complex tasks, sampling from the previous two models (DrIOP) improved performance. For example, in Section 4.4.2 using sampling from two models, we were able to generate all six substructures; however, using just the previous model, we only generated the first five.

We postulate that this is due to the specialisation of the agent at each step. For substructure generation, the success of each generation attempt is dependent on the model randomly generating a structure that maximises the reward. If, after an RL step, the current agent has learned to produce a very narrow selection of structures, the likelihood that it will randomly step out of this local minimum and produce a molecule that satisfies the next reward function is low. Therefore, over-specification at any step is harmful in the long run, and for the best performance the model should learn to generate a diverse set of molecules that satisfy the reward function. Figure 8 shows the pairwise tanimoto similarity of molecules sampled from the agents taken at each step of SrIOP (Figure 8a), DrIOP (Figure 8b) and ReCL (Figure 8c). The figure shows that for SrIOP and DrIOP, the first step leads to agents that produce diverse sets of molecules (blue) with mean tanimoto similarities ∼ 0.2. However, for SrIOP we observe a significant increase in the similarity between molecules generated at steps 2 to 5. For DrIOP we see more pairs of molecules with low similarities for later steps (e.g., steps 2 and 3). Conversely, for ReCL we observe a gradual positive shift in the tanimoto similarity distributions as each new model becomes slightly more specialised. We believed that this would propagate forward and allow DrIOP to solve more complex reward functions, as shown in Figure 7.

**Figure 8:**
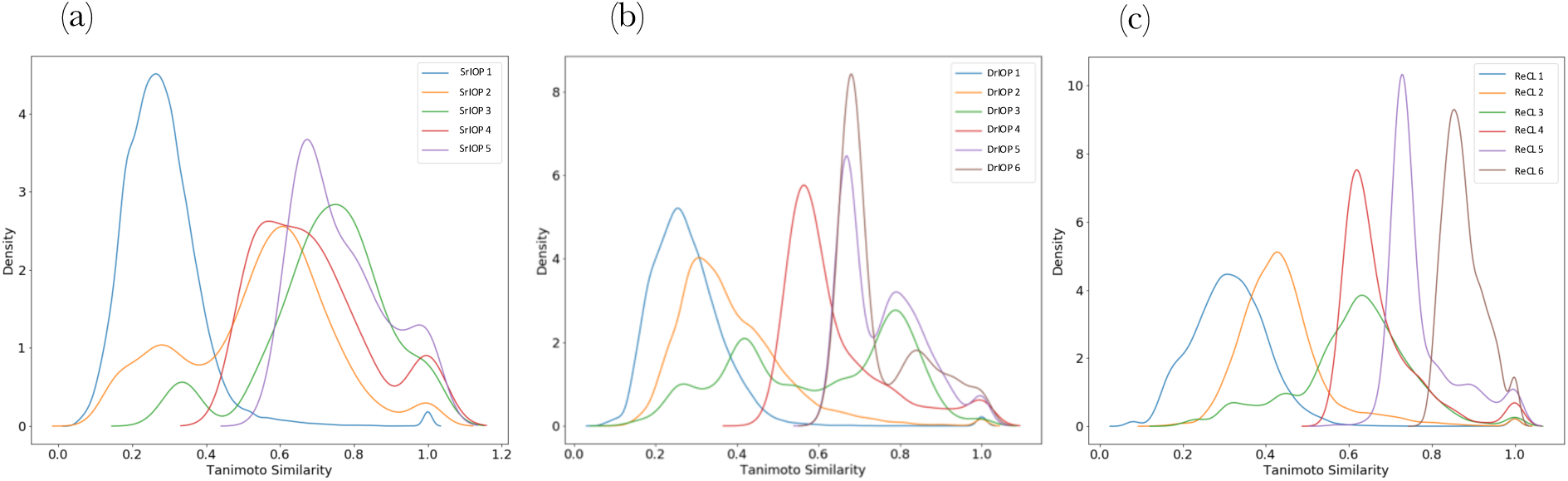
Comparison of model specialisation during (a) SrIOP, (b) DrIOP, and (c) ReCL. Tanimoto similarities of each pair of molecules in the generated set at each step by all methods were calculated. SrIOP shows large increase in similarity for later steps (e. g., 2 to 5) suggesting early specialisation. DrIOP and ReCL have more gradual increase in pairwise tanimoto similarities suggesting more gradual specialisation.

Despite their differences in performance, both implementations of rIOP are useful. The specification we observed with SrIOP allows the model to exploit a chemical property and generate a set of molecules with a very narrow property distribution. In contrast, long optimisation regimes will benefit from the reduced specificity of DrIOP.

### 4.5 Limitations of SMILES-Based Molecular Generators

We have investigated the performance of, and proposed novel, SMILES-based, deep learning, molecular generation tools. These tools learn to generate novel one-dimensional SMILES representations of three-dimensional molecular structures (see methods). SMILES-based tools are popular because they only require simple architectures and can be trained quickly. However, SMILES do present some challenges; namely, they do not detail the three-dimensional structure of a molecule beyond atomic connections and there are several ways to represent the same molecule. Canonical SMILES provide a standard method to generate SMILES; however, it has been shown that SMILES-based models trained on random SMILES show improved model coverage and a reduction in overfitting.^37^

The lack of structural information and inherent redundancy in SMILES can cause SMILES-based models to struggle to fully understand the chemical and structural relationships between molecules. This is because the similarity between two SMILES strings is not well correlated with the similarity between the chemical structures they represent. This limitation of SMILES-based methods and its effects can be seen in our study. For example, when we tried to generate molecules with increasingly complex structures, the performance of the model was heavily dependent on the strings used to represent each substructure. We found that for the best performance, the difference between each string representing a new target structure should be minimised.

To examine the limitations of SMILES-based molecular generators, we attempted to generate molecules that included a target substructure using multiple different SMILES strings to represent the intermediate substructures. We used ReCL and SrIOP to generate molecules with a series of increasingly complex substructures. For each molecule, we enumerated five alternate SMILES for each target intermediate substructure. We found that the choice of SMILES directly affects the performance of the models. Around a quarter of all substructure generation attempts failed (no molecules were generated with the final target substructure) across all molecules (Table 4). But, for each failed attempt, an alternative series of SMILES representing the same molecules was successful. Figure 9 shows the total number of SMILES sampled from the final agent that included the desired substructure and the total number of distinct scaffolds present in successful attempts for two structurally similar molecules (A and B). For molecule A, both ReCL and SrIOP fail to generate the final target substructure at least once, and both methods fail in the final step (supplementary information). For molecule B, we were able to generate substructures regardless of the choice of SMILES even though the intermediate and final substructures were almost identical compared to those of molecule A. This further highlights the issues caused by SMILES as you would expect similar performance across both models given the targets’ structural similarity. Instead, the SMILES used has the largest effect on model performance.

**Table 4:** Proportion of failed substructure generation attempts using different SMILES across all molecules tested using ReCL and SrIOP.

| Method | Number of attempts | Number of failed attempts | Percentage of failed attempts / % |
| --- | --- | --- | --- |
| **ReCL** | 40 | 8 | 20 |
| **SrIOP** | 40 | 12 | 30 |
| **Total** | 80 | 20 | 25 |

**Figure 9:**
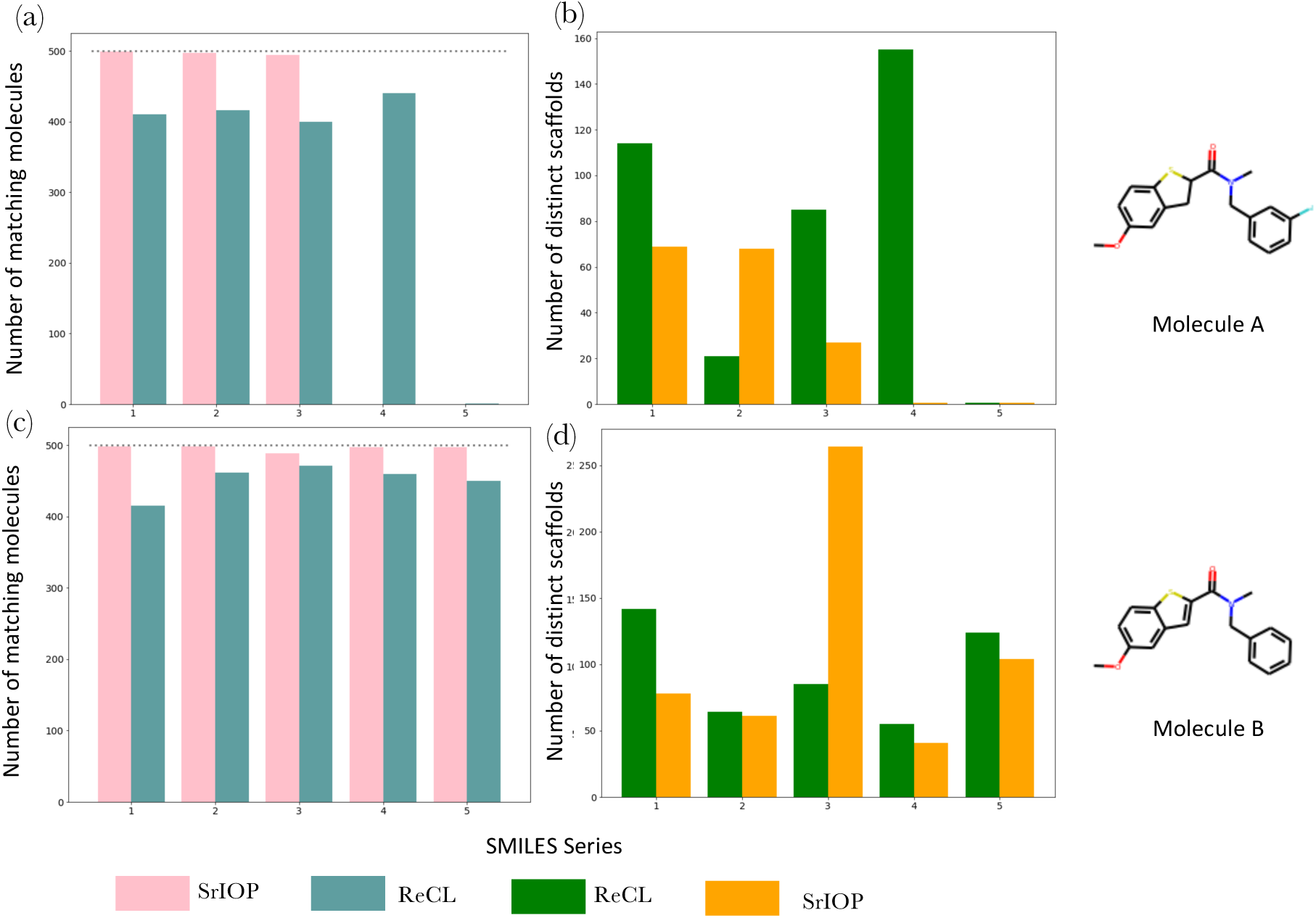
Effects of SMILES choice on substructure generation performance using ReCL and SrIOP. For each molecule 5 sets of different intermediate SMILES were generated, then used during optimisation. Each SMILES variation at each step represented the same molecule. (a) and (c) are the total number of SMILES sampled from the final agent that included the target substructure (500 molecules were sampled). SrIOP (pink) and ReCL (blue). (b) and (d) are the total number of distinct scaffolds present in the successful samples. SrIOP (orange) and ReCL (green). (a) and (b) correspond to molecule A, while (c) and (d) correspond to molecule B. The figure shows that the SMILES choice directly affects optimisation performance and diversity of generated molecules. For example, optimisation of molecule A failed (no molecules matching the desired substructure) at least once with both methods, despite intermediate SMILES at each step across all sets representing the same structure.

The choice of SMILES also has a large effect on the diversity of the generated molecules. For molecule B using SrIOP series 3 (Figure 9c) we generated more than 250 distinct scaffolds across the 500 molecules sampled, while all other SMILES series generated between 50 and 150 scaffolds with ReCL and SrIOP. Similar fluctuations in performance were observed for both ReCL and SrIOP across all molecules tested (supplementary information).

Molecular representations such as SELFIES^38^ and Deep-SMILES^39^ attempt to overcome some of the issues of SMILES in machine learning. However, higher dimensional representations that include structural information are likely to be a more powerful way to represent molecules.

## 5 Conclusions

We have investigated how well deep RL molecular design methods can search beyond the chemical space represented in training data and the effects of the composition of the training data on generated molecular sets. The results show that it is possible to control the distribution of molecules generated by altering the reward function. However, we demonstrate how standard methods (REINVENT) can fail towards the edge of the training data distribution. We found that it is possible to generate molecules with properties that are not present in the training data; nevertheless, we showed that the representation of the desired molecular profile affects the distribution of the generated molecular library. We highlight the lack of control standard methods provide in terms of composition, particularly diversity, of generated molecules and the limitations of SMILES-based molecular generation methods. To overcome some of these issues, we propose a new curriculum learning approach, recurrent iterative optimisation procedure (rIOP), to help boost the diversity of generated molecules when few or no examples of the desired molecular profile are present in the training data. Using this method, we generate structures similar to a series of unseen target structures and outperform other curriculum learning approaches (ReCL). We describe SrIOP and DrIOP, which enable a user to control the diversity of generated molecules for simple and complex optimisation tasks. Using several SMILES representations of the same molecule when generating target structures, we show how the choice of SMILES directly affects the success and performance of SMILES-based tools. Therefore, our method, like any method based on SMILES or other one-dimensional representations will be hampered by the lack of direct structural information.

## 6 Code & Data Availability

The raw data that supports the findings of this study are available from the corresponding author upon request.

The code used in this study is available at https://github.com/m-mokaya/riop.git

## 7 Acknowledgement

This work was supported by funding from the Engineering and Physical Sciences Research Council (EPSRC) and Exscientia.

## Supplementary Information

### S1. Reinforcement Learning Reward Profiles

We used simple step reward functions in all our experiments. For each property, we selected a range of values which would return a positive reward, and then all SMILES generated within that range returned a value of one. All SMILES outside the reward range returned zero. For non-numerical optimisation tasks (for example, substructure generation), if optimisation was successful, the model returned a value of one; otherwise, zero was returned. Figure S1 shows two example reward functions. Figure S1a shows the profile of a reward range; all values within this range received a reward of one. Figure S1b shows a reward profile with a limit where any value greater than 3 received a reward of one. Table 1 shows the reward profiles for all our property optimisation experiments.

**Figure S1:**
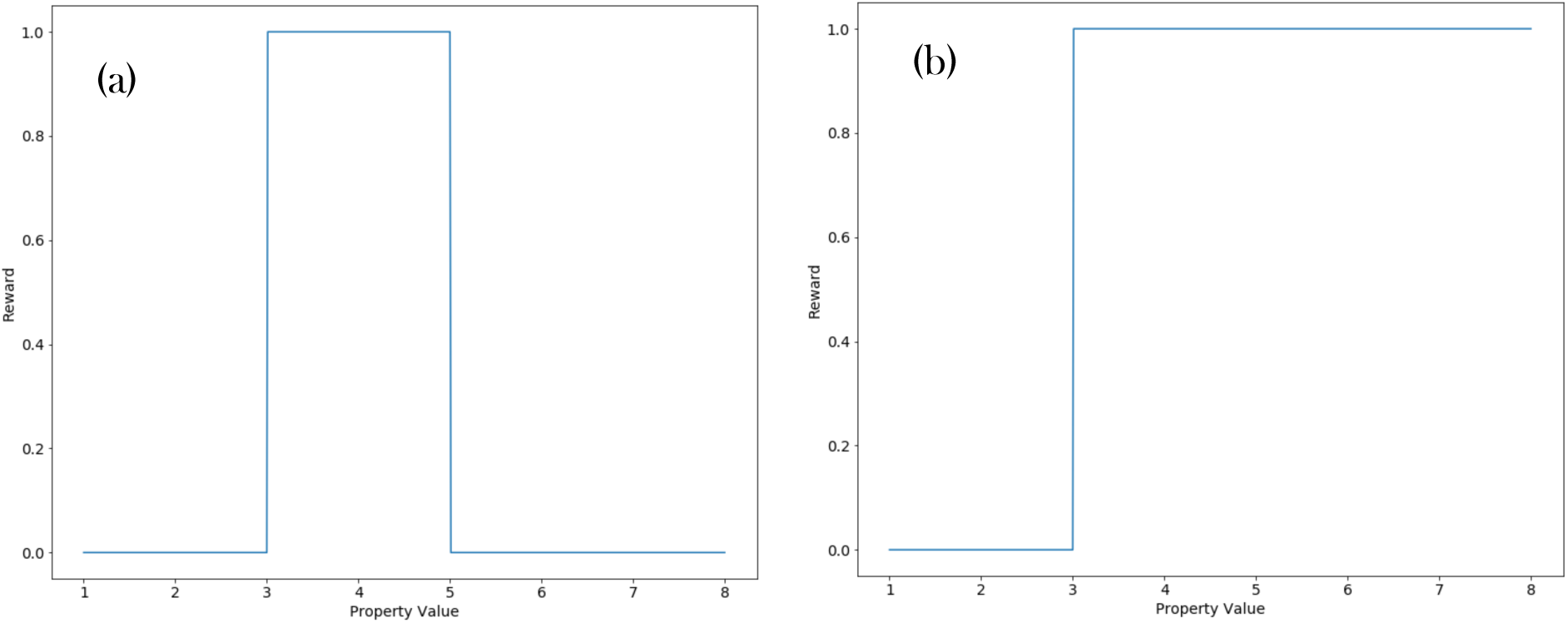
Example reward functions used in our study. (a) reward profile for experiment with a reward range. All molecules inside the range return a reward of one, all outside return zero. (b) reward profile for experiment with reward threshold. All molecules greater than the threshold return one, all other SMILES return zero. For a limit, the same shape would be observed, instead all SMILES less than the limit would return one, all other smiles would return zero.

### S2. Control of generated libraries

Our results show how we can often control the property distribution of generated molecular sets by changing the reward profile during reinforcement learning. We show this to be possible between property-specific values; however, outside of these values, optimisation fails. We demonstrate this with LogP and hydrogen-bond acceptor (HBA) count optimisation. Figure S2 shows the distribution of all optimisation attempts for HBA count between 0 and 350 HBA’s.

**Figure S2:**
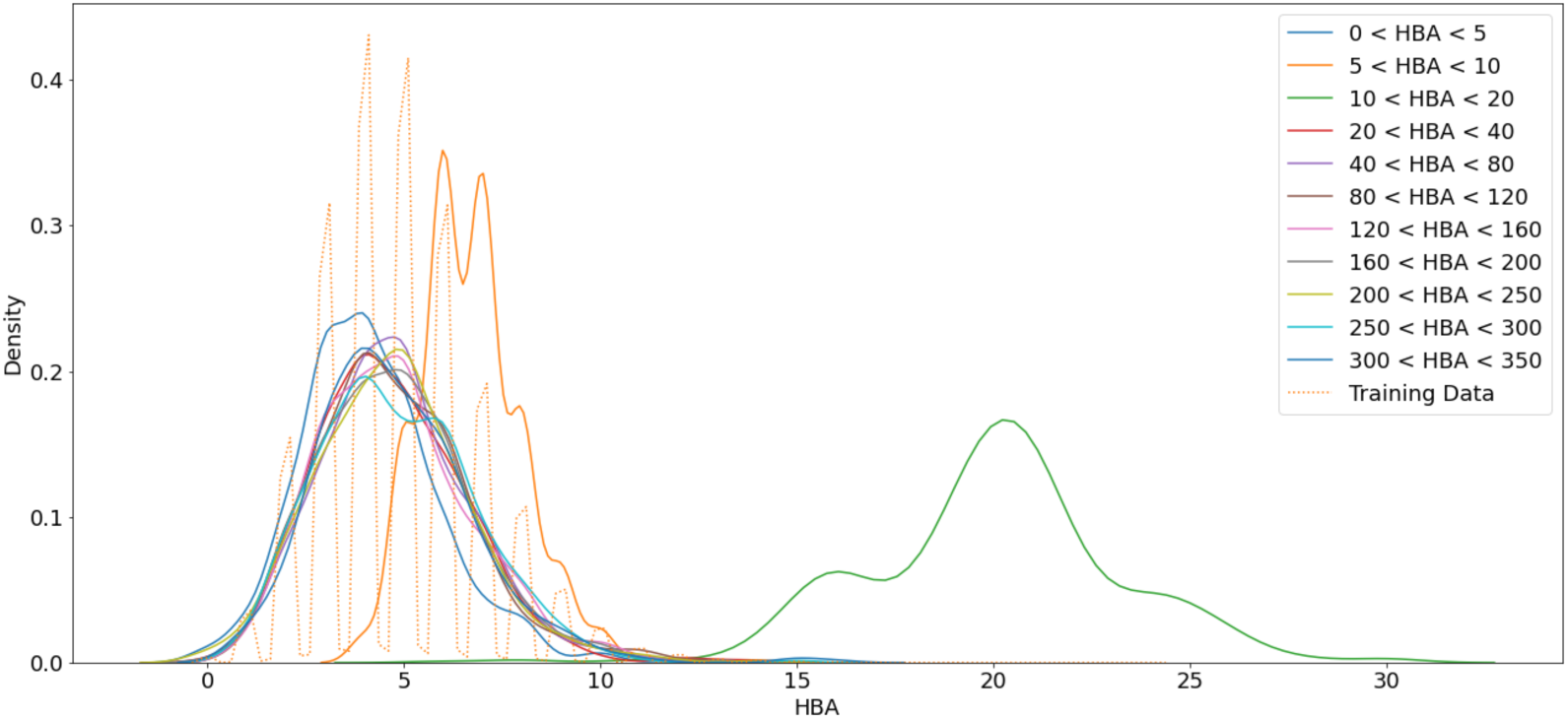
Distribution of generated set for HBA count optimisation. Each line corresponds to the property distribution of the molecules sampled from an agent trained with a reward range detailed in the legend. The distribution of training data (dotted) is also provided.

Figure S3 shows a series of molecules each generated with increasing LogP reward profiles. We see a gradual increase in the LogP of each molecule as the reward range increases; however, at the low extreme (−15 < LogP < -10) optimisation fails and a molecule similar to the training data (LogP between 0 and 10) is produced. Similar results are seen in Figure S2 during hydrogen-bond (HBA) count optimisation. We saw gradual increases in the HBA count until about. 20, beyond this optimisation fails and molecules similar to the training data are produced.

**Figure S3:**
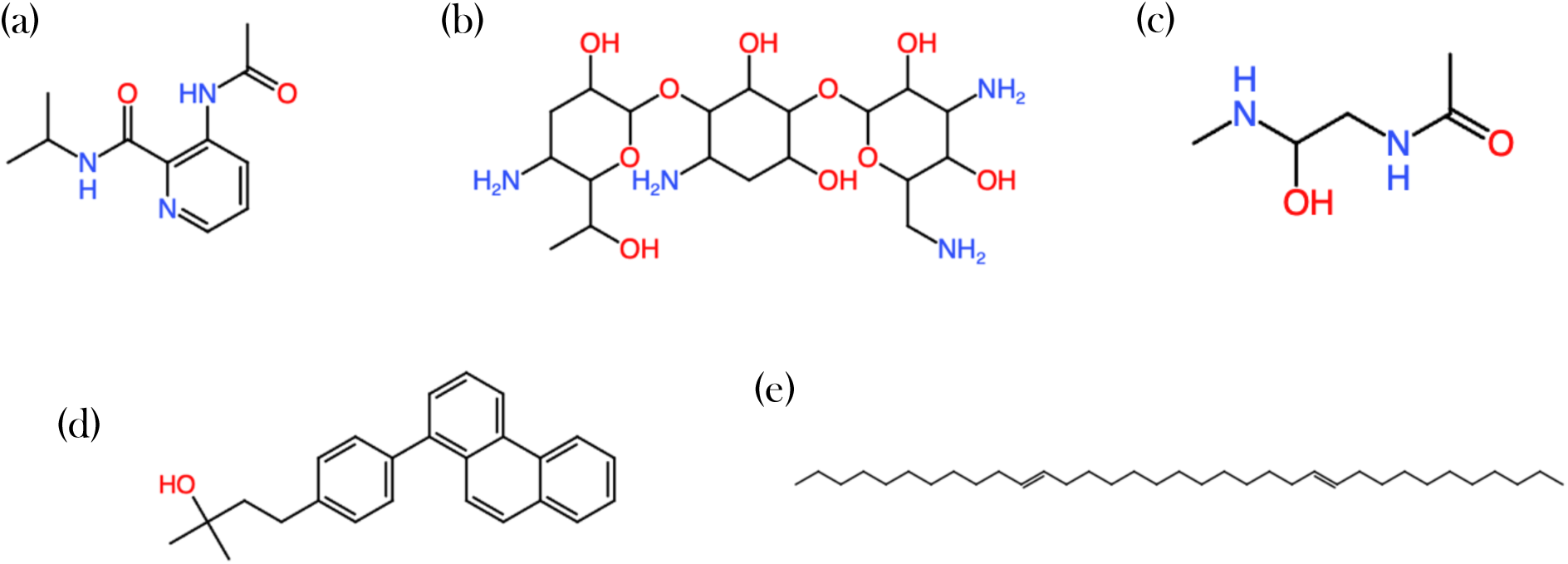
Example molecules from LogP optimisation. Each molecule was sampled from an agent optimised to produce molecules with LogP between (a) - 15 and - 10, (b) – 10 and - 5, (c) - 5 and 0, (d) 5 and 10, (e) 10 and 15.

**Figure S4:**
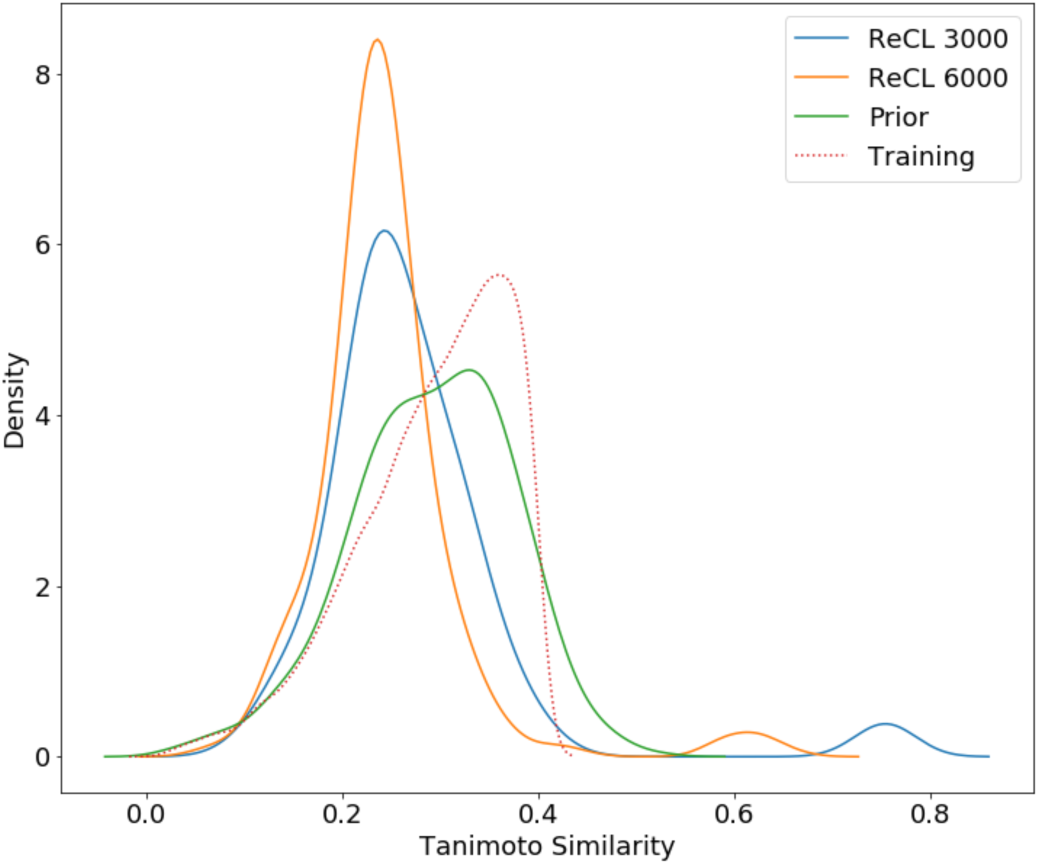
Distributions of generated sets of molecules from ReCL for similarity generation task. Agent training was done for 3000 (blue) and 6000 (orange) (with) iterations to better match SrIOP training regime. Training (dotted) and prior model (green) distributions are provided for reference.

In our comparison of SrIOP to a similar curriculum learning method (ReCL) we tested how well both methods are able to generate molecules similar to target structure with no similar molecules in training data. In our results, we show that ReCL struggles with a complex target structure (Figure 5b), while SrIOP is able to generate similar molecules (Figure 6). One key difference between the methods is the total number of iterations allowed during training. Under the standard protocol, ReCL is assigned a total of 1500 training iterations for all intermediate tasks. The agent will only move on to the next task when it has satisfied the previous one. On the contrary, SrIOP trains for exactly 1000 iterations per intermediate task. This means that for the same set of intermediate tasks, SrIOP is able to train for more iterations. To ensure that any difference in performance is not a result of training iterations, we increased the total number of iterations allowed for ReCL from 1500 to 3000 and 6000 (Figure S4). We found that this had no effect on the results as both models are unable to generate molecules with a tanimoto similarity greater than 0.8 to the target structure.

### S3. Effects of percentage representation

In all our experiments all training datasets were ∼800,000 molecules, and contained either 0%, 2%, 5% 7% or 10% of the molecules within the reward profile. Table S1 detailed the reward thresholds for each property in this experiment. Valid molecules with a property greater than the threshold received a reward of one. All other molecules returned zero reward.

**Table S1:** Reward thresholds and dataset sizes for percentage representation experiments.

| Property | Dataset Size | Reward Threshold |
| --- | --- | --- |
| Hydrogen Bond Acceptors | 798,471 | 8 |
| Molecular Weight | 799,248 | 527 |
| TPSA | 799,063 | 122 |
| QED | 797,569 | 0.83 |

**Figure S5:**
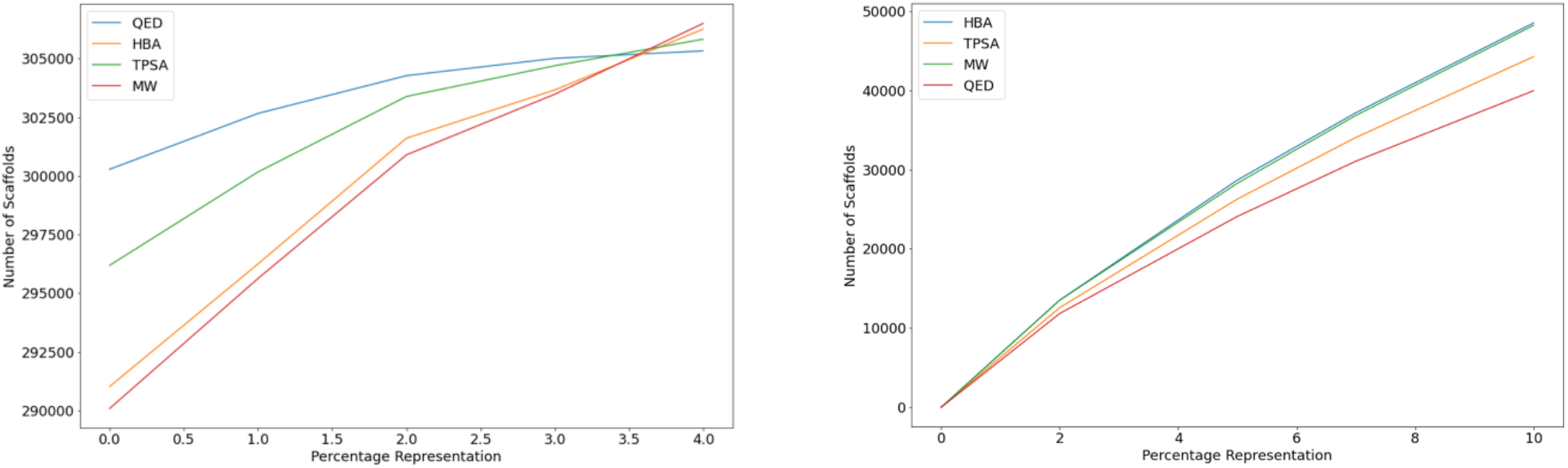
(a) Number of scaffolds present in each training dataset for each curated dataset across four properties. QED (blue), HBA (orange), TPSA (green), molecular weight (red). (b) Number of scaffolds in the reward range region of the same datasets.

**Figure S6:**
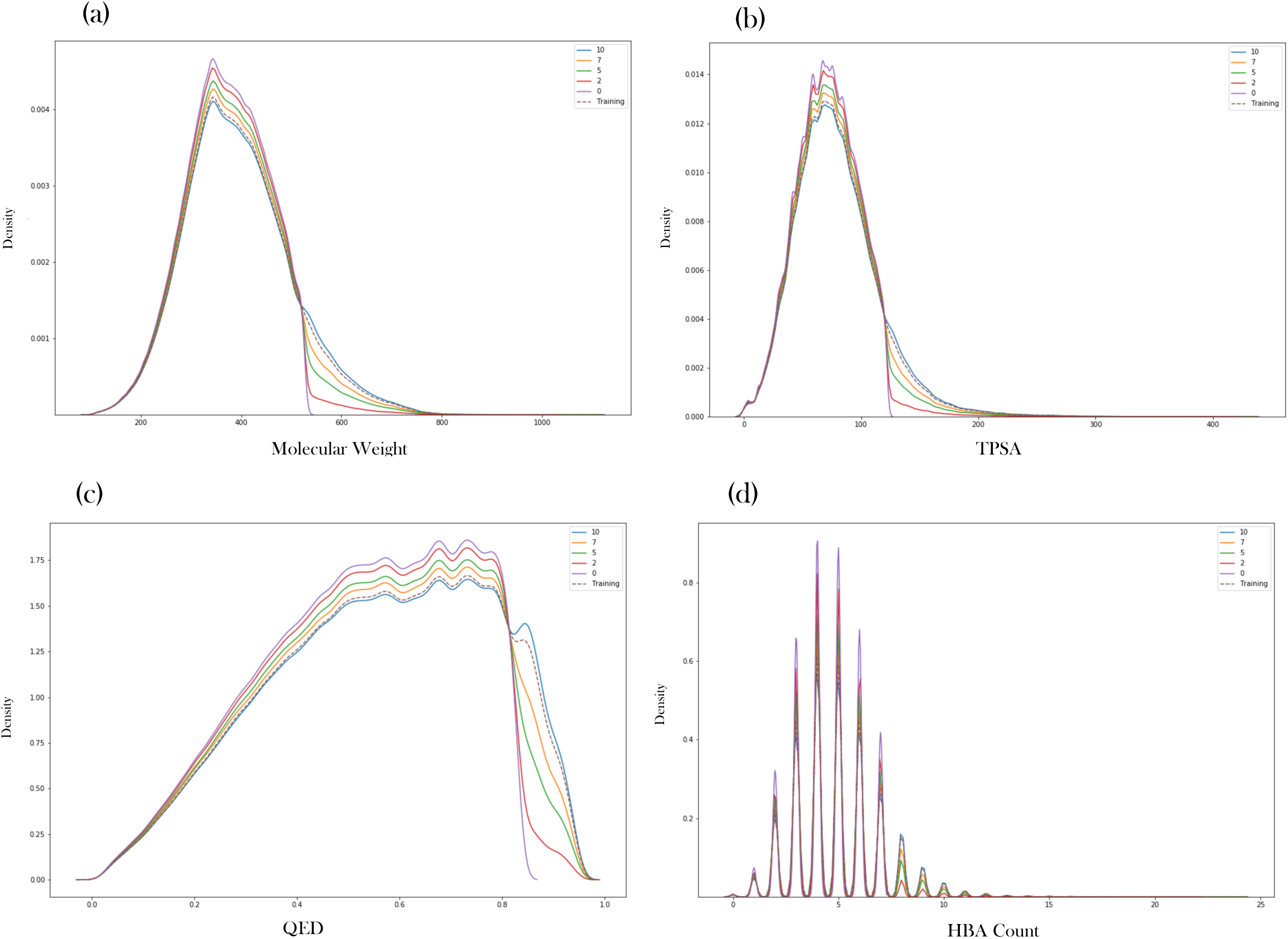
Training data distributions for four properties (a) Molecular weight, (bb) TPSA, (c) QED and (d) HBA count, and percentage representations tested.

**Table 2:**
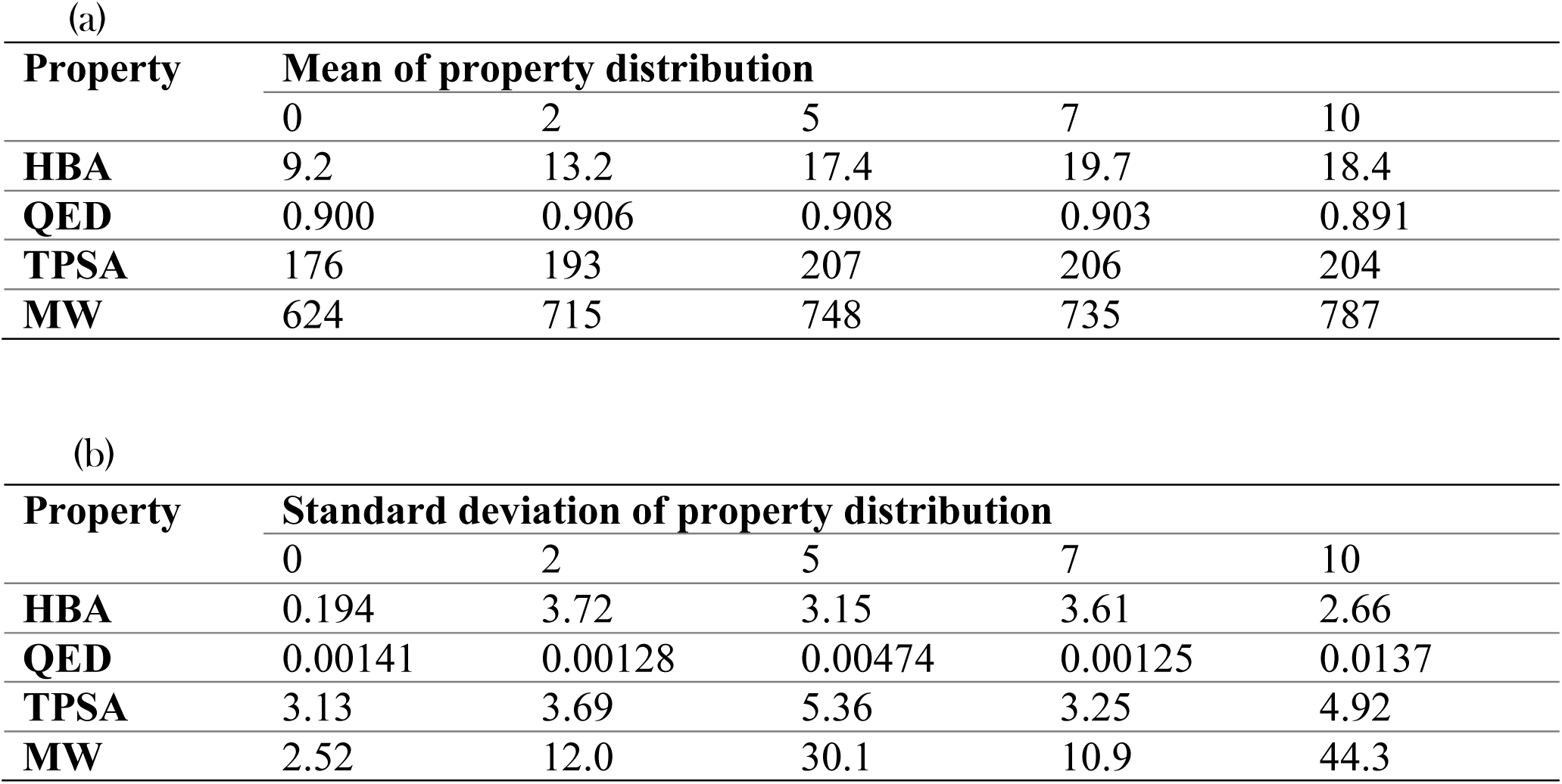
(a) Mean and (b) standard deviation of generated datasets property distributions for three properties at five different training data representations. The values are the average taken from three repetitions.

**Figure S7:**
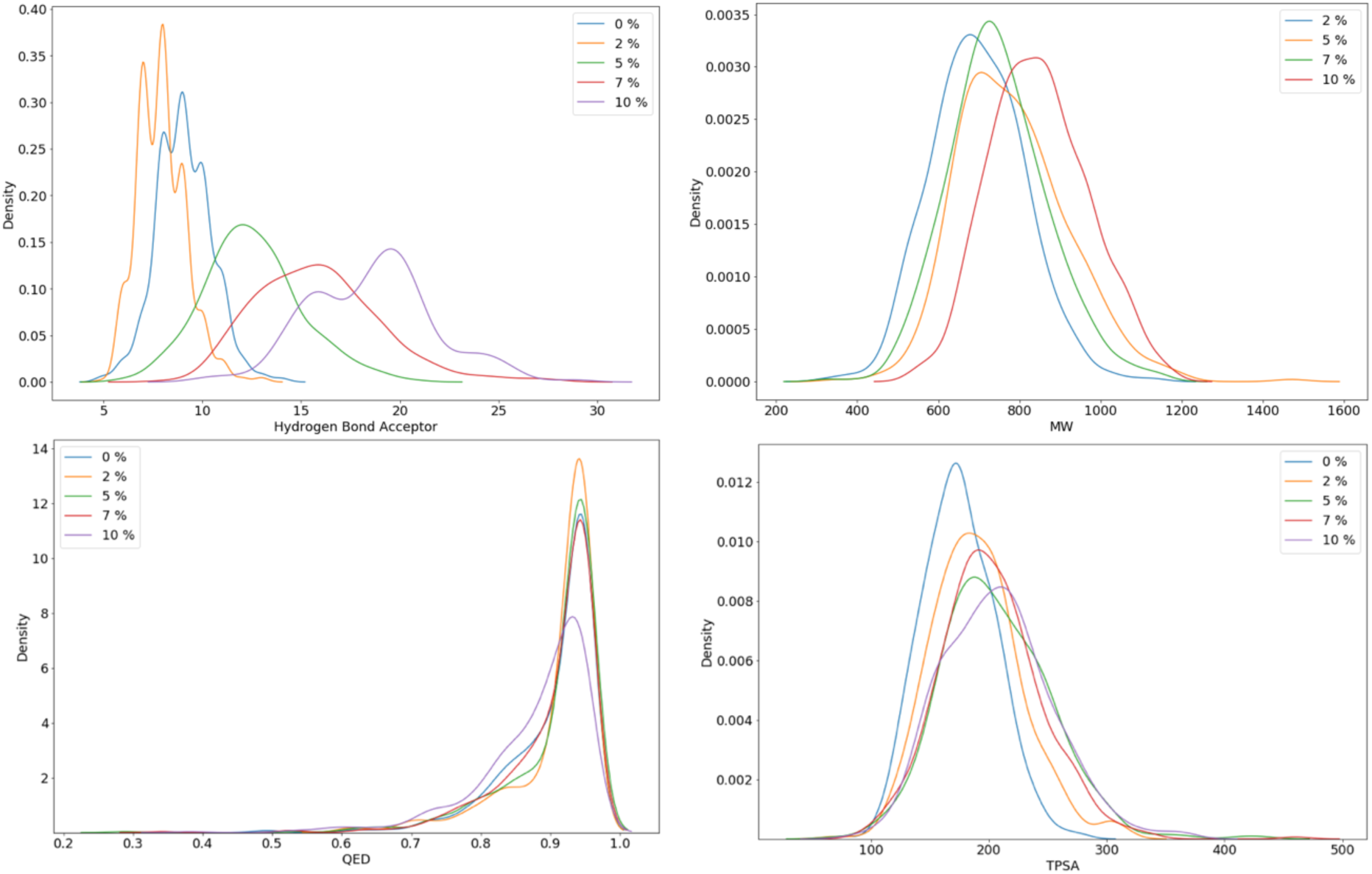
Property distributions for four properties, (a) HBA, (B) molecular weight, (c) QED, (d) TPSA, that where the same optimisation regime was attempted for each using different training data. Each property was tested with 0% (blue), 2% (yellow), 5% (green), 7% (red) and 10% (purple) of the training data matching the reward profile. As the training representation increases, we see a shift in the property distribution. Suggesting that the percentage representation does affect the composition of generated sets of molecules.

### S5. Limitations of SMILES

We show how SMILES choice directly affects the performance of molecular generators. We show that single model rIOP is unable to generate molecules that include a substructure given a series of the increasingly difficult substructures (Figure 7). shows the series of SMILES used for both ReCL and rIOP. ReCL is able to generate all 6 substructures, whereas single model rIOP can only generate 3. We do show how double model rIOP can also generate all six substructures. We suspected that altering the SMILES used would affect the performance of single model rIOP and found alternative routes (figure B) that allowed more of the substructures to be generated. Although we did not find a route to the final structure, we believe that an exhaustive search of possible SMILES combinations would lead to successful substructure generation.

**Figure S8:**
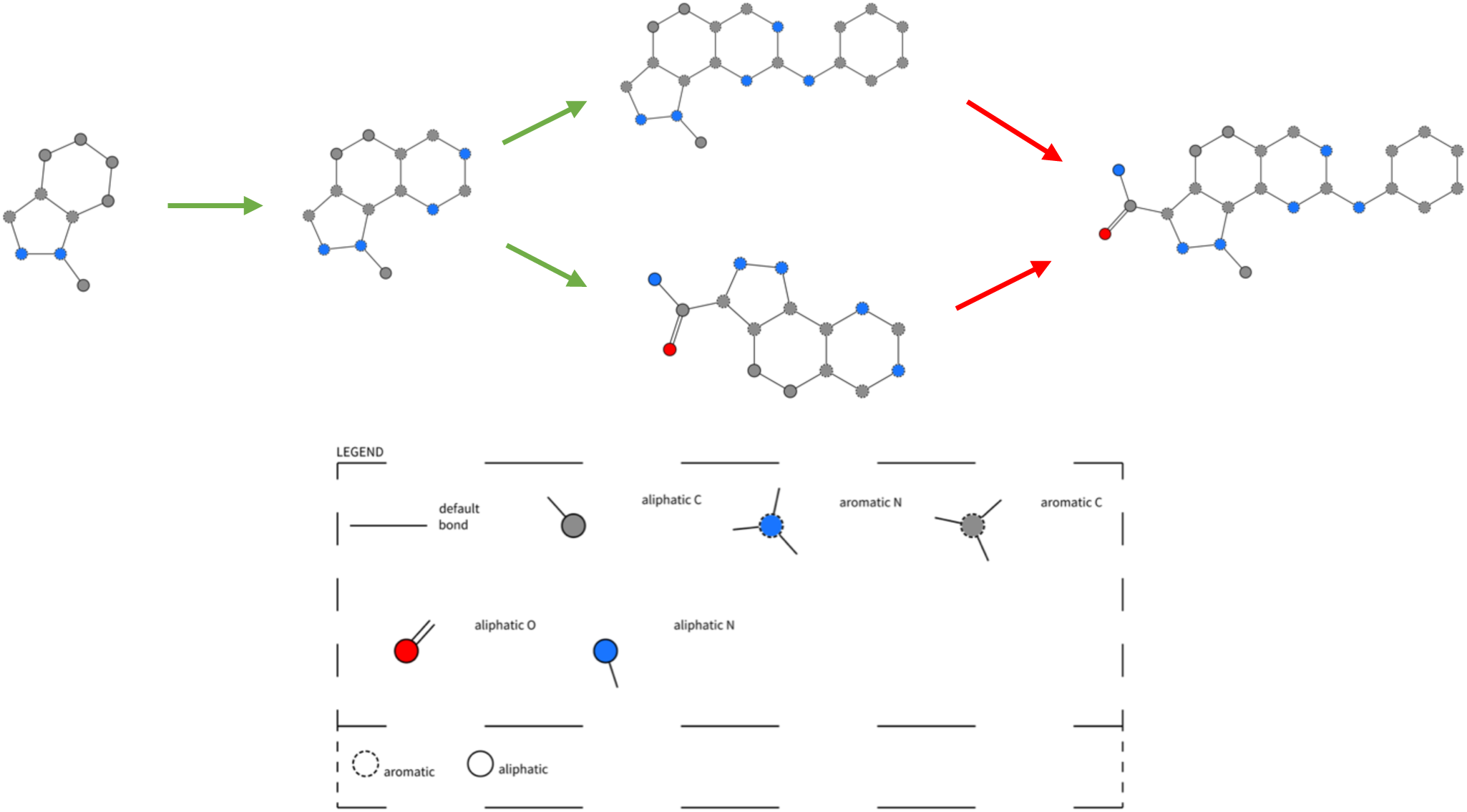
Diagrams showing our various routes used in attempts to generate molecules that included the final substructure. Green arrows represent successful steps where the molecule following the arrow was included in generated molecules. Red arrows are unsuccessful steps.

We have shown the dependence of molecular generators on SMILES choice. To further this, we conducted an experiment on several different molecules from Chembl and curated a series of target substructures. Figure S9 shows the molecules used in our experiments.

**Figure S9:**
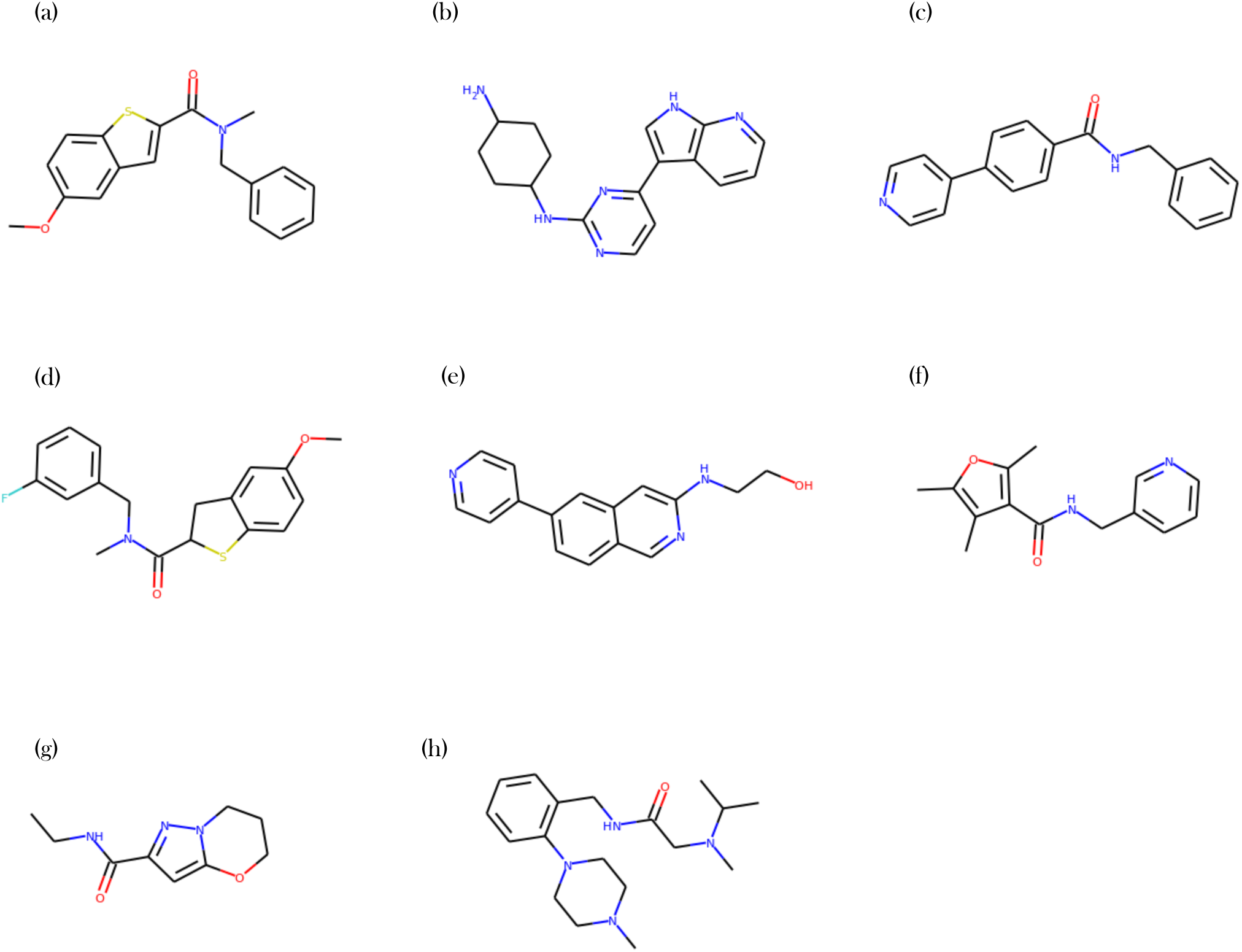
Diagram of all molecules used to demonstrate the effects of SMILES choice on model performance.

**Figure S10:**
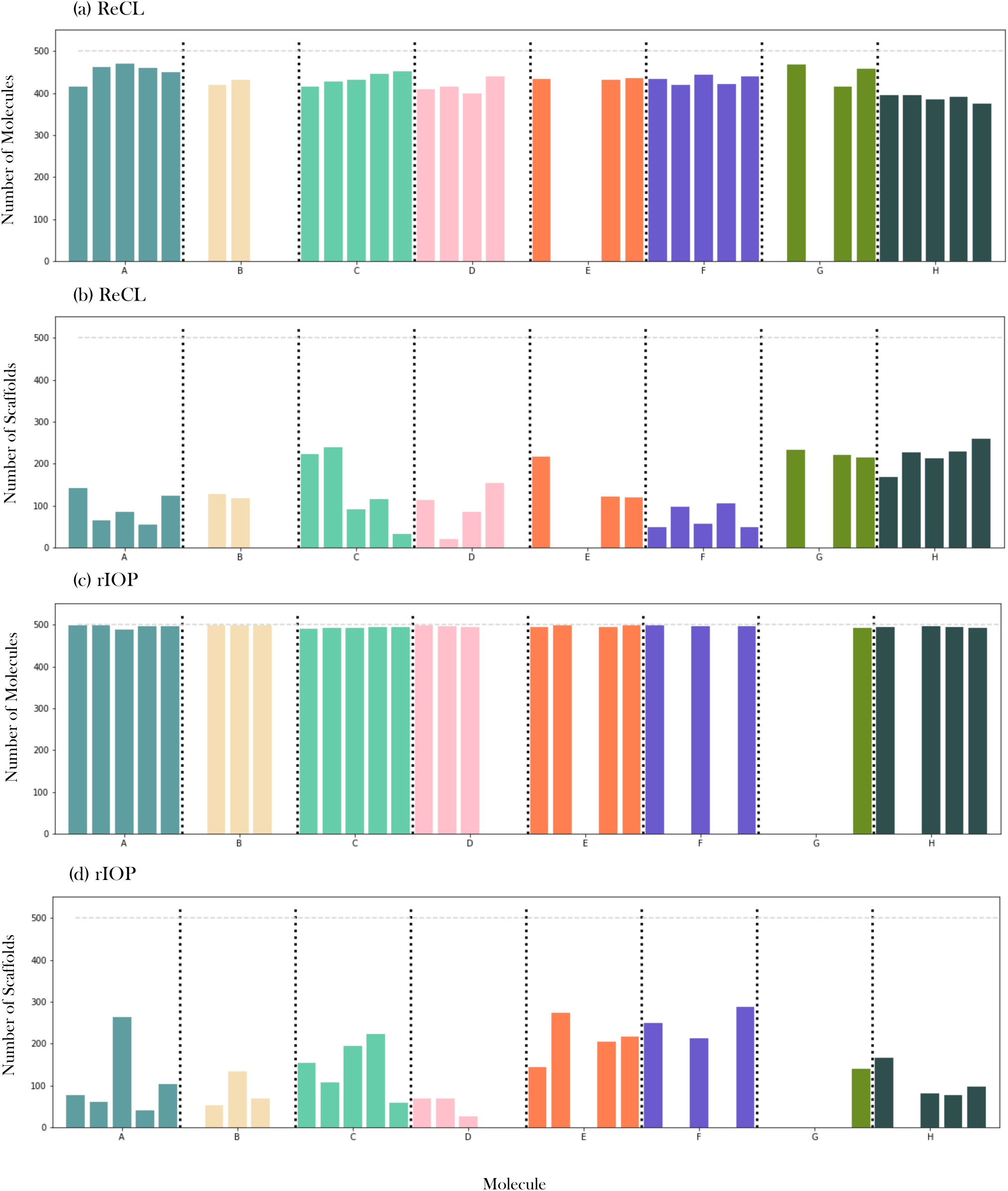
Effect of SMILES on substructure optimisation performance using ReCL (a and b) and single model rIOP (c and d). For each molecule (A-H), 5 sets of intermediate SMILES were enumerated and used during optimisation. In (a) and (c) the coloured bars represent the total number of molecules generated by the final agent (500 molecules were sampled) that included the desired substructure (Figure S9). In (b) and (d), the bars show the total number of distinct scaffolds in the same sample. The figure shows that the SMILES choice directly affects optimisation performance and diversity of generated molecules. For example, optimisation of molecule B failed (no molecules matching the desired substructure) three times using ReCL (a) and twice using rIOP (c) despite intermediate SMILES at each step across all sets representing the same structure.

**Figure S11:**
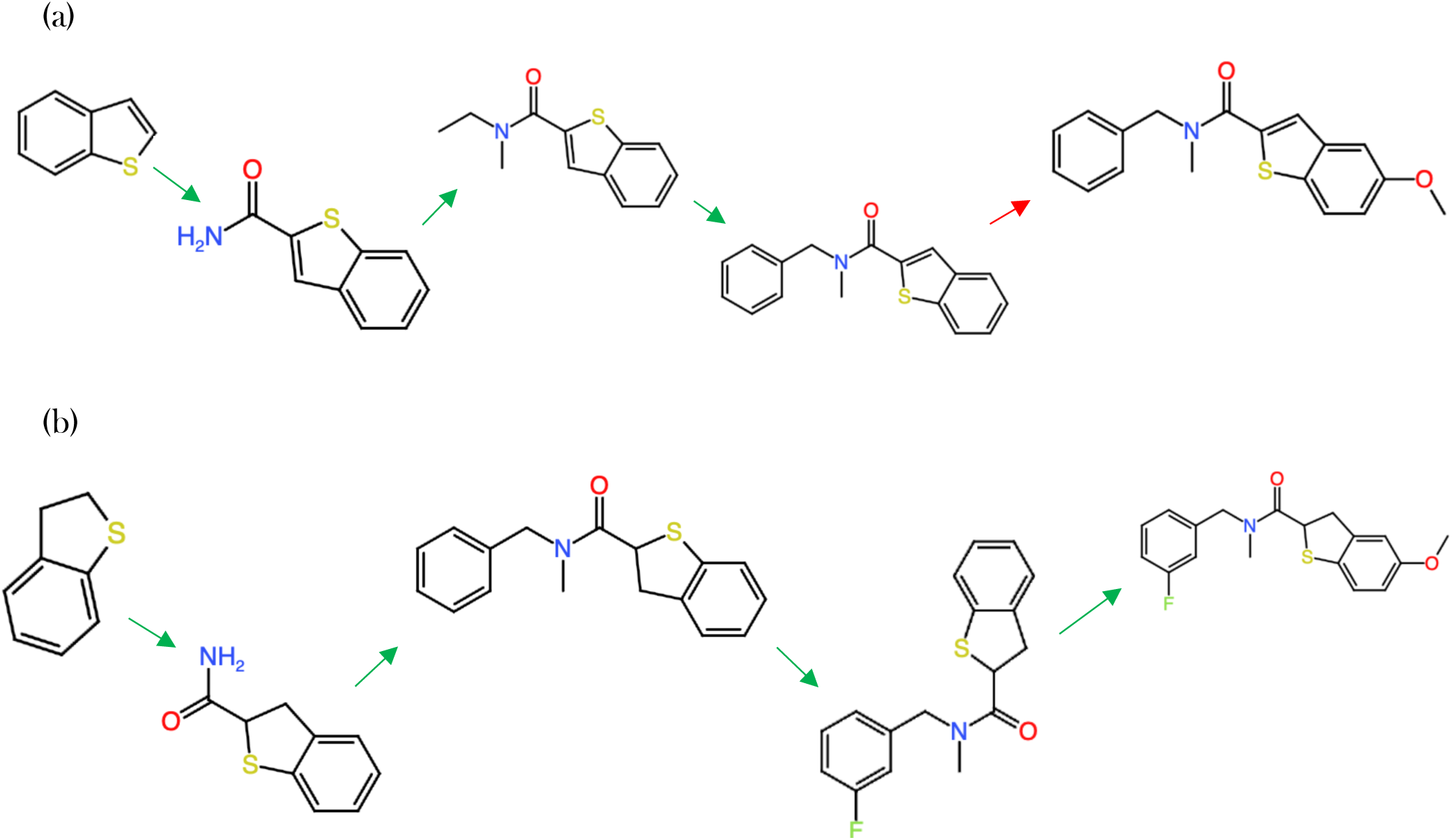
Diagram displaying the intermediate and final substructures we aimed to generate for two molecules. (a) Molecule A and (b) Molecule D from Figure S9. The substructure following a red arrow was not successfully generated after optimisation. Substructures following a green arrow were generated. Despite the intermediates used for both molecules being very similar, both methods tested (ReCL and SrIOP fail on the last step for molecule A.

**Table 3:** Number of molecules that include the target substructure and number of distinct scaffolds for each set of SMILES for (a) molecule A and (b) molecule D.

| Run | Molecules with Substructure |  | Number of Scaffolds |  |
| --- | --- | --- | --- | --- |
|  | SrIOP |  | ReCL |  |
| 1 | 498 | 78 | 415 | 142 |
| 2 | 498 | 61 | 462 | 64 |
| 3 | 489 | 264 | 471 | 85 |
| 4 | 497 | 41 | 460 | 55 |
| 5 | 497 | 104 | 450 | 124 |

(b)
| Run | Molecules with Substructure |  | Number of Scaffolds |  |
| --- | --- | --- | --- | --- |
|  | SrIOP |  | ReCL |  |
| 1 | 499 | 69 | 410 | 114 |
| 2 | 497 | 68 | 416 | 21 |
| 3 | 494 | 27 | 400 | 85 |
| 4 | 0 | 0 | 440 | 155 |
| 5 | 0 | 0 | 0 | 0 |

